# Experimental evidence for reproduction to increase mitochondrial respiration, but decrease mitochondrial efficiency, in females of a short-lived bird

**DOI:** 10.1101/2025.05.12.653466

**Authors:** Matteo Beccardi, Pablo Salmón, Oscar Vedder

**Author notes:** Corresponding author: Matteo Beccardi, Address: Institute of Avian Research, An der Vogelwarte 21, D-26386 Wilhelmshaven, Germany. Shared senior authorship (Pablo Salmón, Oscar Vedder). **Author Contributions:** Conceptualization and experimental design: OV, PS and MB. Data collection: MB. Data analysis: MB, with feedback from PS and OV. Writing: MB and OV, with feedback from PS.

## Abstract

Reproduction is generally considered as one of the most energy-demanding processes, causing a trade-off with survival. Although this “cost of reproduction” plays a pivotal role in theory on ageing and life-history evolution, the physiological mechanisms underlying it remain largely unresolved. As mitochondria synthesize 90% of the energy available to eukaryotic cells, their functioning may be crucial in mediating the cost of reproduction. In this study, we therefore experimentally manipulated the reproductive status (gamete production) of male and female Japanese quail (*Coturnix japonica*), and tested for effects on mitochondrial respiration. We found that reproduction led to an overall increase in mitochondrial O_2_ consumption in females. This was primarily driven by a large increase in proton leak, as O_2_ consumption for adenosine triphosphate (ATP) synthesis only increased slightly. Hence, reproductively active females had a severely reduced efficiency in ATP production, compared to non-reproductive females. In males, mitochondrial respiration and its efficiency were unaffected by reproduction, and similar to that in non-reproductive females. We suggest the large increase in proton leak in reproductive females to represent an adaptive mechanism to mitigate the production of reactive oxygen species (ROS), and to be an unavoidable consequence of elevated ATP synthesis. The latter would cause constraints on energy availability for reproductive females, and thereby trade-offs with other energy demanding processes, like somatic maintenance. The absence of effects of reproduction on male mitochondrial respiration confirms the general view that gamete production is less costly for males.

**Significant statement:** Reproduction is one of the most costly biological processes; not only in terms of energy, but also in terms of its effect on survival. Mitochondria, the primary energy sources in eukaryotic cells, may mediate these costs. By experimentally manipulating the reproductive activity of male and female Japanese quail, we demonstrate that female egg production leads to increased mitochondrial oxygen consumption, primarily due to an increase in proton leak, reducing the efficiency in energy production. We suggest the increased proton leak to be an adaptive response to reduce reactive oxygen species production, but to limit the energy available for processes such as somatic maintenance. Male reproduction did not affect mitochondrial function, confirming the general view that sperm production is relatively cheap.

## INTRODUCTION

Reproduction is generally considered to be one of the most energy-demanding processes, with higher reproductive investment compromising future survival – the so-called “cost of reproduction” (1). This cost of reproduction is pivotal in theory on ageing and life-history evolution (1–3), but the physiological mechanisms that underlie it remain largely unresolved (4–6). Mitochondria are likely to be important in mediating the cost of reproduction, as they synthesize most of the energy available to eukaryotic cells, causing their functioning to affect the organism’s energy availability and its allocation between reproduction and other energy-demanding processes, such as somatic maintenance (6–8). Knowledge on how the functionality of mitochondria is influenced by reproduction may therefore be crucial for understanding the physiological mechanisms underlying the cost of reproduction.

Mitochondria are small organelles present in all eukaryotic cells that via oxidative phosphorylation convert adenosine diphosphate (ADP) to adenosine triphosphate (ATP), which is the chemical fuel that supplies more than 90% of the energy demand at the cellular level (9). Extrinsic and intrinsic factors, such as environmental conditions, age, or physiological state, are known to affect the functionality of mitochondria (e.g., 10–15), but whether reproduction *per se* has an effect on mitochondrial functioning has, to our knowledge, *hitherto* not been experimentally tested.

An experimental test for an effect of reproduction should ideally compare reproducing and non-reproducing individuals that do not differ in anything else than reproduction (16). Silencing of gamete production may, however, be difficult to achieve without also manipulating the individual’s resource availability. In this respect, Japanese quail (*Coturnix japonica*) – small short-lived Galliform birds (17) – are ideal, because their gamete production is extremely sensitive to photoperiod (18, 19), and thus relatively easy to manipulate under controlled conditions. In birds, reproduction involves several energy-demanding processes, but among them, female gamete production (i.e., egg laying) is one of the most energetically costly (20, 21). Hence, if the energetic cost of gamete production affects mitochondrial functionality, we expect to see a difference in mitochondrial functioning between egg laying and non-egg laying female Japanese quail. Gamete production in male birds (i.e., sperm production), on the other hand, is generally considered to be much less energetically costly (22), representing one of the defining differences between the sexes across species (23–25). We would therefore expect to observe a much reduced effect of gamete production on mitochondrial functioning in male Japanese quail, if it is specifically the energetic cost of reproduction that affects mitochondrial function.

In this study, we randomly allocated Japanese quail of both sexes to a reproductively active or a reproductively inactive group, by manipulating their gamete production via the photoperiod they were exposed to (16 vs. 8 h of light per day). Because the erythrocytes of birds contain functional mitochondria (26), we assessed mitochondrial function in a minimally-invasive way in small blood samples. Previous research confirmed that measures of mitochondrial function obtained from blood samples are correlated with those of other tissues, and can thus indicate overall mitochondrial functioning (12, 27–29). With the use of a high-resolution respirometry system, and the addition of specific mitochondrial agonists and antagonists, we induced different respiratory states that provide insight into several measures of mitochondrial respiration and efficiency per sample, which we then compared between reproductively active and inactive females and males.

## RESULTS

The experimental manipulation was successful in causing a large difference in egg laying rate between females assigned to the reproductively active and reproductively inactive treatment (active: 0.85 eggs/day; inactive: 0.01 eggs/day; χ^2^ = 52.38, p < 0.001). The probability of male reproductive activity was confirmed to be similarly different between the treatments (active: 1.00; inactive: 0.10; t = -22.05, p < 0.001).

### Females

In females, the endogenous baseline mitochondrial O_2_ consumption (termed ROUTINE) was ca. 9% higher in the reproductively active *versus* inactive group (p < 0.001, Fig. 1, Table S1). However, this was mostly due to reproduction causing a ca. 29% increase in the mitochondrial respiration linked to proton leak (LEAK; p < 0.001, Fig. 1, Table S1), as the mitochondrial respiration linked to ATP synthesis via oxidative phosphorylation (OXPHOS) only increased with ca. 5% (p = 0.088, Fig. 1, Table S1). The maximum capacity of O_2_ consumption of the electron transport chain (ETS) was increased with ca. 19% (p < 0.001, Fig. 1, Table S1).

**Figure 1.**
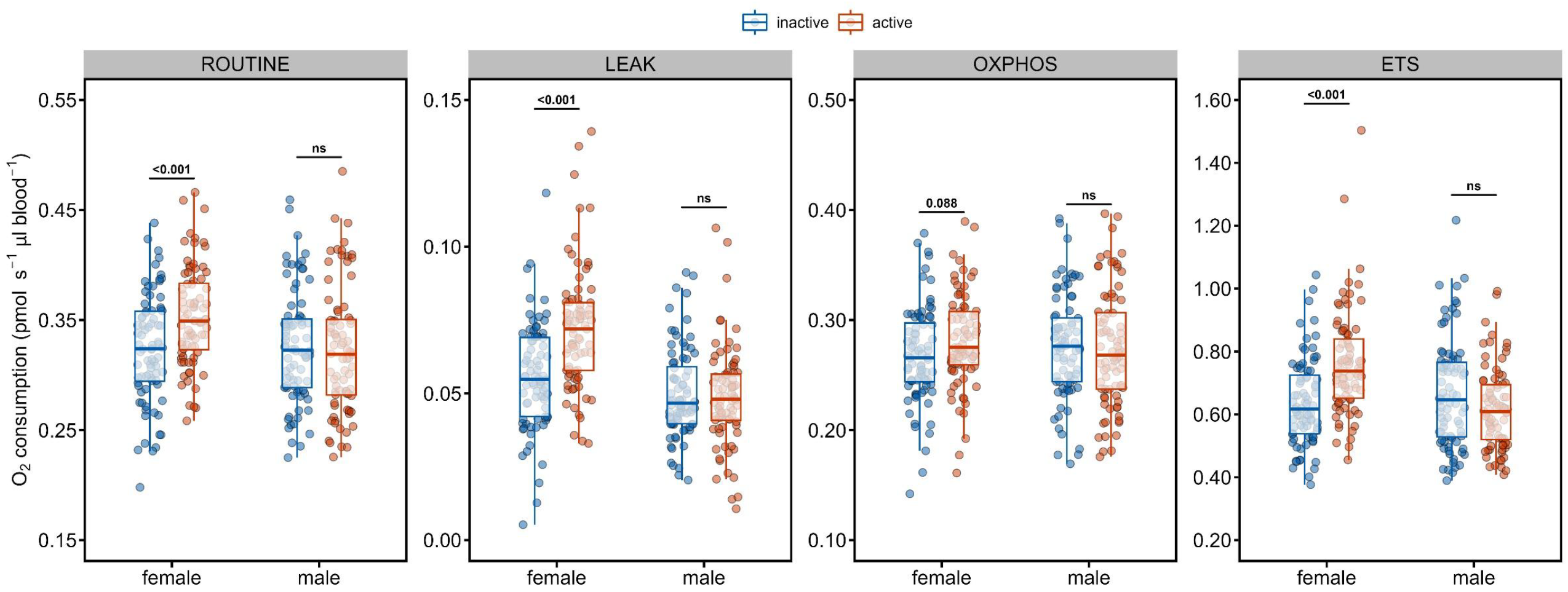
Box-plots, and underlying raw datapoints, showing the effect of reproductive treatment on each mitochondrial respiration trait, in male and female Japanese quail. Reproductively active is expressed in red, while reproductively inactive is expressed in blue. In each box-plot, the thick red/blue line represents the median, while the box encompasses the interquartile range, and the whiskers extend to the most extreme data points within 1.5 × the interquartile range outside the box. Significance of within-sex comparisons are shown in the graphs; further group comparisons are presented in Table S1.

These variable increases in mitochondrial traits led to a reduced efficiency in endogenous respiration for ATP production (R-L control efficiency; p < 0.001, Fig. 2, Table S2), and a lower tightness of the electron transport chain during ATP production when forced to operate at maximum capacity (E-L coupling efficiency; p = 0.031, Fig. 2, Table S2), in the reproductively active females. The proportion of ETS that remained as a surplus under endogenous conditions (E-R control efficiency) was higher in the reproductively active group (p = 0.015, Fig. 2, Table S2).

**Figure 2.**
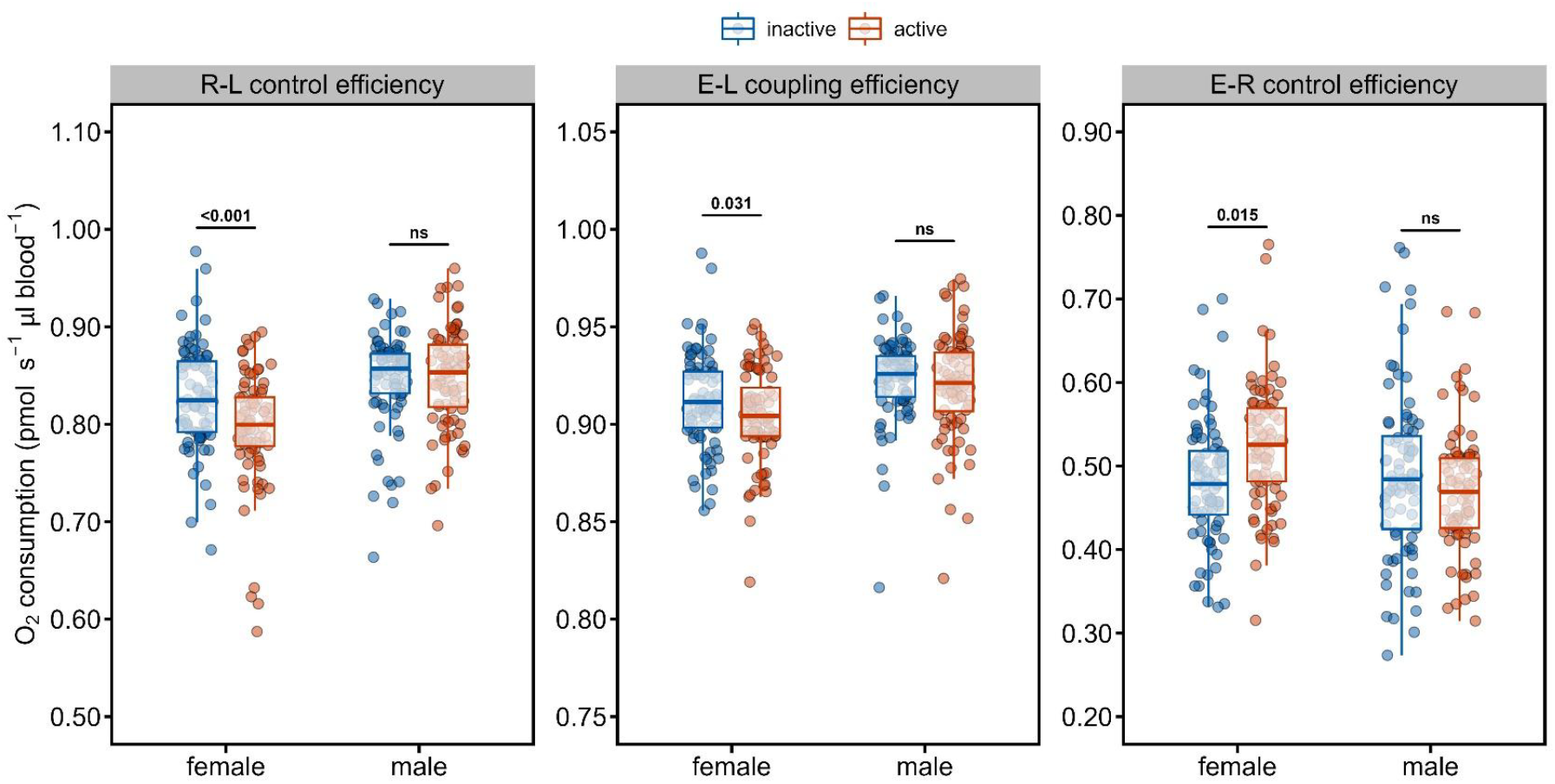
Box-plots, and underlying raw datapoints, showing the effect of reproductive treatment on each mitochondrial efficiency trait, in male and female Japanese quail. Reproductively active is expressed in red, while reproductively inactive is expressed in blue. In each box-plot, the thick blue/red line represents the median, while the box encompasses the interquartile range, and the whiskers extend to the most extreme data points within 1.5 × the interquartile range outside the box. Significance of within-sex comparisons are shown in the graphs; further group comparisons are presented in Table S2.

### Males and sex differences

In males, there was no evidence for a difference between the reproductively active and inactive group for any of the traits quantifying mitochondrial respiration (all p > 0.100, Fig. 1, Table S1), or efficiency (all p > 0.239, Fig. 2, Table S2), and trait levels were not different from those of reproductively inactive females (all p > 0.109, Fig 1, Table S1). Consequently, reproductively active males had significantly lower values for ROUTINE (p < 0.001, Fig. 1, Table S1), LEAK (p < 0.001, Fig. 1, Table S1), and ETS (p < 0.001; Fig. 1, Table S1), but not OXPHOS (p = 0.302; Fig. 1, Table S1), compared to reproductively active females.

Between the sexes, in the reproductively active group, females had lower values for the R-L control efficiency (p < 0.001, Fig. 2, Table S2) and the E-L coupling efficiency (p < 0.001, Fig. 2, Table S2), but higher values for the E-R control efficiency (p = 0.002, Fig. 2, Table S2). On the contrary, in the reproductively inactive group, only the E-L coupling efficiency differed between the sexes (p = 0.013, Fig. 2, Table S2), with lower values in females.

## DISCUSSION

By experimentally manipulating the reproductive status of Japanese quail, we showed that reproduction affects mitochondrial respiration and efficiency in females, but not males. Specifically, we found that reproductively active (i.e., egg laying) females had a disproportionate increase in proton leak and maximum capacity of the electron transport chain, compared to their increase in O_2_ consumption linked to ATP production via oxidative phosphorylation. This led to a reduced efficiency in endogenous respiration linked to ATP production, and a lower tightness of the electron transport chain when forced to operate at maximum capacity. Reproductively active females also maintained a higher proportion of the electron transport chain as a surplus under endogenous conditions, and therefore utilised less of their maximum capacity for O_2_ consumption, compared to reproductively inactive females.

Considering that egg production is an energetically expensive processes (20–22), a reduced efficiency in O_2_ consumption for ATP production in egg-laying females may appear counterintuitive and potentially maladaptive. A previous study in birds that compared mitochondrial function of incubating and chick-rearing pied flycatchers (*Ficedula hypoleuca*) interpreted higher O_2_ consumption linked to proton leak during incubation as a potential functional response to a higher need for the heat dissipation that is associated with proton leak (11). Yet, domesticated Japanese quail have lost the tendency to incubate their eggs (30) and we never observed any incubation behaviour, making this an unlikely explanation in our case. Instead, the “uncoupling to survive” hypothesis posits that the seemingly wasteful mitochondrial proton leak may act as an adaptation to protect against excessive production of reactive oxygen species (ROS) (8). According to the “free radical theory of ageing”, ROS cause damage to mitochondrial and nuclear DNA, ultimately leading to an impaired functioning and ageing (31–33). Hence, given that increased oxidative phosphorylation is associated with increased ROS production (34, 35), the disproportionate increase in proton leak may have acted to mitigate such damage, and thereby acted as a mechanism to reduce the cost of reproduction. As such, our results provide an explanation for why reviews and meta-analyses do not consistently suggest higher levels of oxidative stress in reproducing individuals, despite its high energetic cost (36–38).

Nevertheless, the energetic cost of egg laying does need to be met, and in this respect it is surprising that the O_2_ consumption that is linked to ATP production only increased with ca. 5% in egg-laying females. In its basic form, the “disposable soma” hypothesis suggests that energy is the currency that is directly responsible for the trade-off between reproduction and somatic maintenance, and thereby causes ageing (2, 39). Hence, if the increase in O_2_ consumption for ATP production does not cover the energetic cost of egg-laying completely, this could well come at a cost to somatic maintenance, as ATP is also the principal energy source for maintenance processes such as DNA, RNA and protein synthesis (40). Regardless of its exact cause, if increased ATP production is intrinsically linked to a lower efficiency in ATP production due to a disproportionate rise in proton leak, this may ultimately constrain females in their available energy. Individuals could then either economize on reproduction, or on maintenance. For example, a study on zebra finches (*Taeniopygia guttata*) showed that an experimental increase in proton leak, relative to oxidative phosphorylation, reduced egg production (41), while a study in growing Japanese quail showed that increased proton leak was associated with higher levels of DNA damage (12). This may also explain why the reproductively active females utilised less of their maximum capacity for O_2_ consumption – diminishing returns could make it unprofitable.

None of the effects of reproduction on mitochondrial respiration and efficiency that we observed in females were observed in males. Regardless of their reproductive activity, males resembled reproductively inactive females in all mitochondrial traits. Our results thereby confirm that the generally much higher cost of gamete production for females (22), which forms the foundation for sex differences throughout the tree of life (23–25), is reflected in measurements of metabolism at the cellular level. Male costs of reproduction are more likely to result from increased expenditure on competition for territories or matings (24, 42), which, due to pairwise experimental housing, were not present in our study. This further supports our conclusion that it is specifically the energetic cost of egg laying that is responsible for the observed difference in mitochondrial functioning between reproductively active and inactive females. Overall, our findings highlight the crucial role of mitochondrial functioning in mediating the cost of reproduction, and provide an explanation for why trade-offs between reproduction and maintenance processes can even be observed when individuals are not limited in food availability (16).

## MATERIAL AND METHODS

### Experimental design and sample collection

The experiment was carried out with individuals from a captive population of Japanese quail maintained at the Institute of Avian Research in Wilhelmshaven, Germany, in the local winter of 2023 to 2024. Individuals aged between ca. 0.5 and 1.5-years-old, of both sexes (n = 80 males and 80 females), were randomly allocated to two experimental groups. One group of quail was stimulated to become reproductively active in winter, by keeping the individuals in two outdoor aviaries (one per sex) exposed to an artificial photoperiod of 16 h of light per day. The other group was kept in similar aviaries (also one per sex), but exposed to the natural photoperiod, causing no development of reproductive activity in winter.

After ca. 1-2 months of manipulating their reproductive activity in these aviaries, the individuals were transferred to indoor cages in two main rounds (n = 80 individuals in October-December 2023, and 80 individuals in February-April 2024). In these cages (122 x 50 x 50 cm), individuals were kept in pairs (male and female), all with artificial lighting, at ca. 16-18 °C, and with *ad libitum* water and food – a standard layer diet with 19.0% protein, 4.6% fat and 4.8% calcium, and a caloric value of 9.8 MJ/Kg (GoldDott, DERBY Spezialfutter GmbH, Münster, Germany). In both rounds, 20 pairs of the reproductively active group received a 16 h photoperiod throughout sample collection, while 20 pairs of the reproductively inactive group received a photoperiod of 8 h. All pair combinations were randomly selected within the round and treatment group, but avoiding brother-sister pairs. To spread the workload for sample analysis, the transfers from outdoor to indoor were performed with 4 pairs per day (two active and two inactive). All individuals were sampled for mitochondrial function analysis twice: 7 days and 21 days after moving to the indoor cages (n = 320 total samples). In both sampling events, a small blood sample (ca. 100 μL) was collected by puncturing the brachial vein. We always sampled two complete pairs in the morning (one active and one inactive) and two complete pairs in the afternoon, to minimize the storage time of blood before analysis (range: 0.5 - 6 h, at 4 °C). During the entire duration of the indoor sampling period, the reproductive activity of the females was monitored by daily checks for newly laid eggs, which were collected at first appearance. Male reproductive activity was assessed at the blood-sampling events by recording the presence of foam from the proctodeal gland. In Japanese quail, the foam acts as a type of seminal fluid and previous research has shown that its presence is indicative of sperm production (43, 44). All individuals were returned to outdoor aviaries the day after their second blood sample was collected.

### Mitochondrial respiration in whole blood

An Oxygraph-2k high resolution respirometer (Oroboros Instruments, Innsbruck, Austria) was used to analyse mitochondrial respiration in all collected blood samples, following the method described by (45). We analysed 8 samples per day in 4 different runs, selecting the run and respirometry chamber (A or B) randomly per sample. Samples were gently resuspended with a pipette and 50 μL of whole blood were added to 1.95 mL of respirometry buffer ((MiR05; 0.5 mM EGTA, 3 mM MgCl2, 60 mM K-lactobionate, 20 mM taurine, 10 mM KH2PO4, 20 mM Hepes, 110 mM sucrose, free fatty acid bovine serum albumin (1 g l-1), pH 7.1); (46)), previously incubated at 40 °C (12) in the respirometry chamber. Following the re-equilibration of the oxygen levels (after at least 2 min), the chamber was closed and the respiration rate was quantified as the rate of decline in O_2_ concentration in the chamber. Once the O_2_ consumption signal stabilized (after approximately 25-30 min), the basal respiration on endogenous substrates (“ROUTINE”) was measured over a recording period of at least 2 min. We subsequently added 1 μL of 5 μmol L-1 of oligomycin (Merck, Germany) to inhibit the mitochondrial ATP synthesis and record the mitochondrial respiration linked to mitochondrial proton leak (“LEAK”). Volumes of 0.5 μL increments of 1 μmol L-1 of the mitochondrial uncoupler FCCP (Thermo Fisher, Germany) were added until the maximum O_2_ consumption was reached (1.5-2 μL of FCCP), thus recording the artificially induced maximum capacity of the electron transport chain (“ETS”). Finally, the residual or non-mitochondrial O_2_ consumption was quantified by adding 1 μL of 5 μmol L-1 of antimycin A (Merck, Germany) and subsequently subtracted from all respiration states. The mitochondrial respiration linked to ATP synthesis (“OXPHOS”) was calculated by subtracting LEAK from ROUTINE.

In addition, three mitochondrial efficiency parameters were calculated following (46): R-L control efficiency ((ROUTINE - LEAK) / ROUTINE), which is the proportion of endogenous respiration used for ATP production via oxidative phosphorylation. E-L coupling efficiency (1 - (LEAK / ETS)), which is the proportion of ETS maximum capacity linked to proton leak, indicating the tightness of electron transport during ATP production when forced to operate at maximum capacity. E-R control efficiency (1 - (ROUTINE / ETS)), which is measured as the proportion of the maximum working capacity remaining as a surplus under the endogenous cellular conditions.

### Statistical analyses

To test if the experimental manipulation was successful in causing a difference in egg laying rate between females assigned to the reproductively active and inactive treatment, we performed a generalized linear mixed model (GLMM; function “lmer()” from R-package “lme4”) with a binomial error distribution and logit-link function. Whether or not a female laid an egg (no = 0, yes = 1) on each day of the experiment (21 days per female) was included as the dependent variable, while “reproductive treatment” (2 levels: active/inactive) and “female identity” were included as a fixed and random effect, respectively. Given the lower number of repeated observations per individual for males, and the different distribution of data between the two reproductive treatments, the treatment effect on sexual activity during the indoor period in males was tested with a linear mixed model (LMM; function “lmer()” from R-package “lme4”), with “presence of foam” (no = 0, yes = 1) as the dependent variable, “treatment” as fixed effect and “male identity” as a random effect.

To test for effects of the reproductive manipulation on mitochondrial respiration and efficiency parameters, we ran seven separate LMMs, one with each mitochondrial trait as the dependent variable (ROUTINE, LEAK, OXPHOS, ETS, R-L control efficiency, E-L coupling efficiency, E-R control efficiency). In each model, we included “reproductive treatment” (2 levels: active/inactive) and “sex” (2 levels: male/female), and their interaction, as fixed effects. In addition, “sampling day” (2 levels: day 7/day 21), “round” (2 levels: first/second) and “chamber” (2 levels: A/B) were included as fixed effects to correct for potential differences between sampling days or rounds, and O_2_ consumption between chambers, respectively. Finally, “individual identity” and “pair identity” were included as random effects, to account for the two repeated measures per individual, and the potential for cage location or other effects at the pair level, respectively. A normal error distribution was used in each of these models, as a visual inspection of the distribution of the residuals did not reveal marked deviations from normality (function “simulateResiduals()”, from R-package “DHARMa”). We used F-tests based on Satterthwaite approximation for the denominator degrees of freedom to estimate the significance of parameter estimates. The interactions between “reproductive treatment” and “sex” were further explored using pairwise planned comparisons adjusted by Tukey HSD (function “emmeans” from R-package “emmeans”).

All analyses were performed using R software (version 4.3.1, R Foundation for Statistical Computing).

## Acknowledgments and funding sources

We thank Adolf Völk for the quail husbandry, and Sandra Bouwhuis for constructive comments on the manuscript. The study was funded by grant number 428800869 from the German Research Foundation (DFG, Deutsche Forschungsgemeinschaft) to OV. Permission for all experimental procedures involving the quail was granted by the Lower Saxony State Office for Consumer Protection and Food Safety (LAVES), Germany (permit nr. 33.19-42502-04-22-00212).

