## Supplementary material for "Experimental evidence for reproduction to increase mitochondrial respiration, but decrease mitochondrial efficiency, in females of a short-lived bird": Table S1

**-Supporting information-**

**Matteo Beccardi, Pablo Salmón, Oscar Vedder**

### Tables

**Table S1** Summaries of LMMs testing for effects of the reproductive treatment and sex on the mitochondrial respiration variables.

**ROUTINE**

| *Fixed effects and covariates* | *Estimate* | *S.E.* | *df* | *F-value* | *p-value* |
|---|---|---|---|---|---|
| Reproductive treatment (active) | 0.029 | 0.007 | 1 | 5.298 | 0.024 |
| Sex (male) | 0.001 | 0.007 | 1 | 7.075 | 0.009 |
| Sampling day (day 21) | 0.014 | 0.004 | 1 | 12.675 | <0.001 |
| Round (second) | 0.050 | 0.005 | 1 | 79.584 | <0.001 |
| Chamber (B) | 0.005 | 0.004 | 1 | 1.246 | 0.265 |
| Reproductive treatment : Sex | -0.032 | 0.010 | 1 | 8.607 | 0.004 |
| active females - inactive females | 0.029 | 0.007 | | | <0.001 |
| active males - inactive males | -0.003 | 0.007 | | | 0.689 |
| active females - active males | 0.030 | 0.007 | | | <0.001 |
| inactive females - inactive males | -0.001 | 0.007 | | | 0.847 |
| active females - inactive males | 0.027 | 0.007 | | | 0.003 |
| inactive females - active males | 0.001 | 0.007 | | | 0.997 |
| *Random effects* | *Variance* | *S.D.* | | | |
| Individual identity | 5.11x10$^{-4}$ | 0.022 | | | |
| Pair identity | 3.31x10$^{-5}$ | 0.006 | | | |
| Residual | 1.37x10$^{-3}$ | 0.037 | | | |

**LEAK**

| *Fixed effects and covariates* | *Estimate* | *S.E.* | *df* | *F-value* | *p-value* |
|---|---|---|---|---|---|
| Reproductive treatment (active) | 0.017 | 0.003 | 1 | 9.628 | 0.002 |
| Sex (male) | -0.005 | 0.003 | 1 | 39.821 | <0.001 |
| Sampling day (day 21) | -0.001 | 0.001 | 1 | 0.299 | 0.585 |
| Round (second) | 0.001 | 0.002 | 1 | 0.303 | 0.583 |
| Chamber (B) | -0.003 | 0.001 | 1 | 2.415 | 0.121 |
| Reproductive treatment : Sex | -0.018 | 0.004 | 1 | 16.102 | <0.001 |
| active females - inactive females | 0.016 | 0.003 | | | <0.001 |
| active males - inactive males | -0.001 | 0.003 | | | 0.687 |
| active females - active males | 0.023 | 0.003 | | | <0.001 |
| inactive females - inactive males | 0.005 | 0.003 | | | 0.109 |
| active females - inactive males | 0.021 | 0.003 | | | <0.001 |
| inactive females - active males | 0.006 | 0.003 | | | 0.213 |
| *Random effects* | *Variance* | *S.D.* | | | |

| | | |
|---|---|---|
| Individual identity | 1.05x10-4 | 0.010 |
| Pair identity | 2.17x10-5 | 0.004 |
| Residual | 1.93x10-4 | 0.014 |

**OXPHOS**

| *Fixed effects and covariates* | *Estimate* | *S.E.* | *df* | *F-value* | *p-value* |
|---|---|---|---|---|---|
| Reproductive treatment (active) | 0.012 | 0.007 | 1 | 1.079 | 0.300 |
| Sex (male) | 0.006 | 0.007 | 1 | 0.005 | 0.941 |
| Sampling day (day 21) | 0.015 | 0.003 | 1 | 18.027 | <0.001 |
| Round (second) | 0.049 | 0.005 | 1 | 93.664 | <0.001 |
| Chamber (B) | 0.008 | 0.004 | 1 | 3.812 | 0.051 |
| Reproductive treatment : Sex | -0.014 | 0.010 | 1 | 1.935 | 0.166 |
| active females - inactive females | 0.012 | 0.007 | | | 0.088 |
| active males - inactive males | -0.001 | 0.007 | | | 0.802 |
| active females - active males | 0.007 | 0.007 | | | 0.302 |
| inactive females - inactive males | -0.006 | 0.007 | | | 0.355 |
| active females - inactive males | 0.005 | 0.007 | | | 0.860 |
| inactive females - active males | -0.004 | 0.007 | | | 0.904 |
| *Random effects* | *Variance* | *S.D.* | | | |
| Individual identity | $4.76x10^{-4}$ | $2.18x10^{-2}$ | | | |
| Pair identity | $2.36x10^{-11}$ | $4.86x10^{-6}$ | | | |
| Residual | $1.07x10^{-3}$ | $3.28x10^{-2}$ | | | |

**ETS**

| *Fixed effects and covariates* | *Estimate* | *S.E.* | *df* | *F-value* | *p-value* |
|---|---|---|---|---|---|
| Reproductive treatment (active) | 0.123 | 0.029 | 1 | 3.167 | 0.080 |
| Sex (male) | 0.027 | 0.029 | 1 | 8.392 | 0.004 |
| Sampling day (day 21) | 0.026 | 0.010 | 1 | 6.893 | 0.009 |
| Round (second) | 0.100 | 0.021 | 1 | 22.686 | <0.001 |
| Chamber (B) | 0.005 | 0.013 | 1 | 0.120 | 0.730 |
| Reproductive treatment : Sex | -0.172 | 0.040 | 1 | 17.937 | <0.001 |
| active females - inactive females | 0.123 | 0.029 | | | <0.001 |
| active males - inactive males | -0.048 | 0.029 | | | 0.100 |
| active females - active males | 0.145 | 0.029 | | | <0.001 |
| inactive females - inactive males | -0.027 | 0.029 | | | 0.347 |
| active females - inactive males | 0.096 | 0.029 | | | 0.007 |
| inactive females - active males | 0.021 | 0.029 | | | 0.888 |
| *Random effects* | *Variance* | *S.D.* | | | |

| | | |
|---|---|---|
| Individual identity | 1.24x10$^{-2}$ | 0.111 |
| Pair identity | 7.10x10$^{-4}$ | 0.026 |
| Residual | 8.02x10$^{-3}$ | 0.090 |

**Table S2** Summaries of LMMs testing for effects of the reproductive treatment and sex on the mitochondrial efficiency variables.

**R-L control efficiency**
**(Phosphorylating efficiency)**

| *Fixed effects and covariates* | *Estimate* | *S.E.* | *df* | *F-value* | *p-value* |
|---|---|---|---|---|---|
| Reproductive treatment (active) | -0.034 | 0.009 | 1 | 5.217 | 0.025 |
| Sex (male) | 0.015 | 0.009 | 1 | 25.706 | <0.001 |
| Sampling day (day 21) | 0.010 | 0.004 | 1 | 6.027 | 0.015 |
| Round (second) | 0.021 | 0.007 | 1 | 9.626 | 0.002 |
| Chamber (B) | 0.014 | 0.005 | 1 | 7.436 | 0.007 |
| Reproductive treatment : Sex | 0.036 | 0.013 | 1 | 7.616 | 0.007 |
| active females - inactive females | -0.034 | 0.009 | | | <0.001 |
| active males - inactive males | 0.002 | 0.009 | | | 0.792 |
| active females - active males | -0.052 | 0.009 | | | <0.001 |
| inactive females - inactive males | -0.015 | 0.009 | | | 0.107 |
| active females - inactive males | -0.049 | 0.009 | | | <0.001 |
| inactive females - active males | -0.017 | 0.009 | | | 0.249 |
| *Random effects* | *Variance* | *S.D.* | | | |
| Individual identity | 1.03x10$^{-3}$ | 0.032 | | | |
| Pair identity | 7.83x10$^{-5}$ | 0.008 | | | |
| Residual | 1.46x10$^{-3}$ | 0.038 | | | |

**E-L coupling efficiency**
**(Tightness of the electron transport chain)**

| *Fixed effects and covariates* | *Estimate* | *S.E.* | *df* | *F-value* | *p-value* |
|---|---|---|---|---|---|
| Reproductive treatment (active) | -0.008 | 0.004 | 1 | 4.137 | 0.045 |
| Sex (male) | 0.009 | 0.003 | 1 | 21.248 | <0.001 |
| Sampling day (day 21) | 0.004 | 0.002 | 1 | 3.977 | 0.048 |
| Round (second) | 0.011 | 0.002 | 1 | 14.602 | <0.001 |
| Chamber (B) | 0.006 | 0.002 | 1 | 6.492 | 0.011 |
| Reproductive treatment : Sex | 0.005 | 0.005 | 1 | 1.043 | 0.310 |

| | | | | | |
|---|---|---|---|---|---|
| active females - inactive females | -0.008 | 0.004 | | | 0.030 |
| active males - inactive males | -0.003 | 0.004 | | | 0.432 |
| active females - active males | -0.015 | 0.003 | | | <0.001 |
| inactive females - inactive males | -0.009 | 0.003 | | | 0.013 |
| active females - inactive males | -0.019 | 0.004 | | | <0.001 |
| inactive females - active males | -0.006 | 0.004 | | | 0.345 |
| *Random effects* | *Variance* | *S.D.* | | | |
| Individual identity | $9.42x10^{-5}$ | 0.009 | | | |
| Pair identity | $2.20x10^{-5}$ | 0.004 | | | |
| Residual | $4.28x10^{-4}$ | 0.020 | | | |

**E-R control efficiency**
**(Reverse capacity)**

| *Fixed effects and covariates* | *Estimate* | *S.E.* | *df* | *F-value* | *p-value* |
|---|---|---|---|---|---|
| Reproductive treatment (active) | 0.042 | 0.017 | 1 | 0.834 | 0.362 |
| Sex (male) | 0.008 | 0.014 | 1 | 3.634 | 0.058 |
| Sampling day (day 21) | -0.001 | 0.004 | 1 | 0.021 | 0.884 |
| Round (second) | 0.001 | 0.012 | 1 | 0.022 | 0.881 |
| Chamber (B) | -0.001 | 0.006 | 1 | 0.070 | 0.792 |
| Reproductive treatment : Sex | -0.063 | 0.024 | 1 | 6.673 | 0.010 |
| active females - inactive females | 0.042 | 0.017 | | | 0.014 |
| active males - inactive males | -0.020 | 0.017 | | | 0.239 |
| active females - active males | 0.054 | 0.017 | | | 0.002 |
| inactive females - inactive males | -0.008 | 0.017 | | | 0.634 |
| active females - inactive males | 0.034 | 0.017 | | | 0.194 |
| inactive females - active males | 0.012 | 0.017 | | | 0.896 |
| *Random effects* | *Variance* | *S.D.* | | | |
| Individual identity | 0.005 | 0.071 | | | |
| Pair identity | 0.000 | 0.000 | | | |
| Residual | 0.001 | 0.041 | | | |
